# Intracranial electroencephalography reveals effector-independent evidence accumulation dynamics in multiple human brain regions

**DOI:** 10.1101/2023.04.10.536314

**Authors:** Sabina Gherman, Noah Markowitz, Gelana Tostaeva, Elizabeth Espinal, Ashesh D. Mehta, Redmond G. O’Connell, Simon P. Kelly, Stephan Bickel

## Abstract

Neural representations of perceptual decision formation that are abstracted from specific motor requirements have previously been identified in humans using non-invasive electrophysiology, however, it is currently unclear where these originate in the brain. Here, we capitalized on the high spatiotemporal precision of intracranial EEG to localize such abstract decision signals. Presurgical epilepsy patients judged the direction of random-dot stimuli and responded either with a speeded button press (N=23), or vocally, after a randomized delay (N=11). We found a widely distributed motor-independent network of regions where high-frequency activity exhibited key characteristics consistent with evidence accumulation, including a gradual build-up that was modulated by the strength of the sensory evidence, and an amplitude that predicted subjects’ choice accuracy and response time. Our findings offer a new view on the brain networks governing human decision making.

## Introduction

A perceptual decision is the process of choosing among alternatives on the basis of sensory information, such as judging whether a traffic light is red or green. Such decisions are often tied to particular motor plans (e.g., braking if the light is red). This has motivated a large body of non-human primate neurophysiology work to characterize the neural correlates of perceptual decisions in regions of the brain involved in motor selection/planning, i.e., where activity is selective for the action used to express choice. Activity at these sites is consistent with a process of evidence accumulation, building gradually over time at a rate that is proportional with the strength of the sensory information, reaching a fixed threshold just before a response is made, and reliably predicting the animal’s choice and response time^1–6^.

In complex environments however, decisions about sensory events must often be made even when the appropriate movement is not known in advance, or when one is not required immediately. How such abstract (i.e., motor-independent) decisions are enabled by the brain remains unclear. In recent years, a concerted effort has been made to identify abstract decision signals in both human and non-human research. Single unit recordings in the non-human primate have uncovered signals that reflect choice-relevant categories independently of specific motor requirements, across parietal^7–9^, prefrontal^10^, and midbrain structures^11^. However, the limited spatial coverage in these studies precludes a comprehensive spatial mapping of the generators of similar signals. Meanwhile, human neuroimaging work has identified several regions across the prefrontal and parietal cortex where blood-oxygen-level-dependent (BOLD) signals correlate with the strength of sensory evidence irrespective of response modality ^12–16^. However, the low temporal resolution of functional magnetic resonance imaging (MRI) precludes verification that these regions exhibit the expected evidence accumulation dynamics. Additionally, these studies differ in the assumptions they make regarding the expected BOLD profile of a region involved in decision making, and, consequently, there has been considerable variability across studies in the regions identified with this method.

Human electrophysiology has contributed high temporal precision and whole-brain analyses to the characterization of effector-independent decision dynamics. Several studies have been able to decode choice-selective activity for tasks involving stimulus presence/absence^17^, intensity^18^, or identity^19–21^ judgments. However, an approach focused on choice selectivity could overlook areas of the brain where choice alternatives are represented by spatially intertwined or adjacent neuronal populations that are not separable with the relatively low spatial resolution of these techniques, as neuronal selectivity to choice and task properties can be highly heterogeneous even within small regions (e.g., LIP^22^).

Indeed, several scalp EEG studies have highlighted an abstract signature of evidence accumulation over centroparietal electrodes (the centro-parietal positivity or CPP) that undergoes a positive build-up during deliberation regardless of which alternative the cumulative evidence favors and which, therefore, would likely be missed by classifiers decoding choice-selective activity for two-alternative discriminations ^23–26^. The CPP exhibits a unique combination of characteristics that can be leveraged in analyses designed to identify abstract evidence accumulation signals in intracranial recordings. These include a gradual buildup during deliberation, with a slope that scales with evidence strength^24,27^ and predicts response time irrespective of evidence location^28^ or sensory modality^23^. The CPP is also motor-independent, tracing the cumulative evidence when the stimulus-response mapping is unknown at the time of evidence presentation^29^ and even when no overt action is required at all^23^. An additional feature that distinguishes the CPP from the build-to-threshold signals observed in effector-selective areas is that its amplitude at the time of choice scales with accuracy and decreases as a function of reaction time, in contexts where choices must be reported within a strict deadline^25,27,30^.

While CPP-like signals measured on the scalp appear to be in agreement with the existence of abstract representations of cumulative evidence, their generators in the brain are not currently known. Aside from offering a more detailed map of the neural architecture that supports decision making, establishing its generators would be a critical step toward understanding how decisions are represented in the brain. A scalp-recorded signal like the CPP may reflect the summation of activity from potentially multiple sources and thus obscure more complex spatiotemporal dynamics. Invasive recordings from human intracranial EEG could circumvent these limitations, as this method provides millimeter spatial precision, while preserving the temporal resolution of scalp EEG/MEG recordings^31^. Additionally, it offers broader spatial sampling compared to single-unit recordings.

Here, we recorded intracranial EEG (iEEG) in a large cohort (N=23) of pre-surgical epilepsy patients as they performed perceptual discrimination paradigms designed to allow characterization of evidence-dependent buildup signals in terms of their distinct temporal profile and selectivity for, or independence from, evidence location and motor response (Fig 1a). Subjects viewed two simultaneous random dot patches and were asked to report the direction of random-dot motion occurring in either one with a speeded button press. To evaluate potential decision-related signals arising independently of motor requirements, a subset of participants (N=11) also performed a version of the task in which choices were reported vocally after a randomized delay. We identified multiple locations where high-frequency activity dynamics were consistent with abstract evidence accumulation, primarily distributed across prefrontal, parietal, inferior temporal, and insular regions. Specifically, activity began to build up following the presentation of sensory evidence, showed modulation by the quality of the evidence, and correlated with behavioral speed and accuracy. Importantly, it was independent of the low-level properties of the stimulus (spatial location) and of the modality of the motor response used to express choice. Our data provide an extensive mapping of abstract evidence accumulation dynamics across the human brain and a strong guide to future investigations into the neural substrate of abstract decisions.

**Figure 1.**
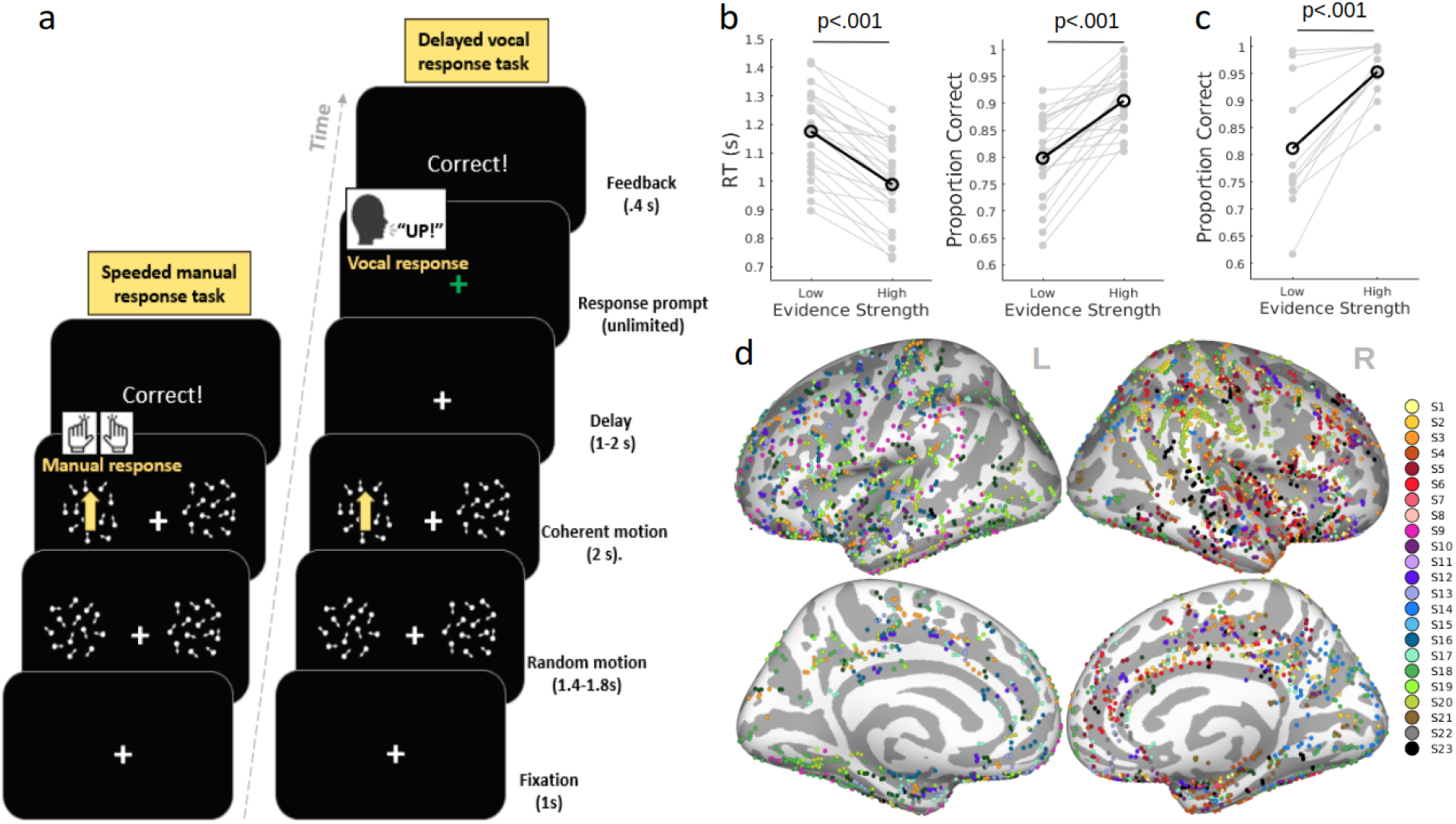
**a.** Behavioral task design. On both tasks, subjects viewed two simultaneous patches of randomly moving dots, located laterally on each side of a fixation cross. After a random delay, motion on one (unpredictable) side became coherent, either upwards (90°) or downwards (270°). In the manual response task (left panel), subjects reported the direction of the coherent motion as quickly and as accurately as possible, with a button click with their left/right hand, whose mapping to direction was counterbalanced across subjects. During the vocal response task (right panel), subjects responded after a random delay, prompted by a visual cue, by verbally naming the direction of motion. Feedback was provided at the end of each trial. **b.** Behavioral performance during the manual response task, as a function of motion coherence (left panel: response time; right panel: choice accuracy). Black data points represent averages across subjects. Gray data points represent individual subject data. **c.** Choice accuracy during the vocal response task as a function of motion coherence. **d.** Intracranial electrode coverage for the manual response task. Each color represents a subject. Electrodes have been projected onto the Freesurfer^32^ average inflated brain (light gray: gyri; dark gray: sulci).

## Results

We recorded iEEG activity while subjects performed a two-stimulus motion discrimination task (Fig 1a). In each trial, two patches of incoherently moving dots appeared simultaneously on the left and right side of a fixation point, and following a random delay, one of them, selected at random, began to move coherently at either a high or low coherence level. The direction (upwards or downwards) of this motion was to be reported by the subject as quickly and as accurately as possible.

### Manual response task

#### Behavior

In the speeded manual response version of the task (Fig. 1a), subjects were instructed to report their choice within a 2000 ms deadline by means of a button press with their left/right hand. As expected, we found that behavioral performance was strongly modulated by the strength of sensory evidence (i.e., motion coherence) across subjects, with stronger evidence leading to improved choice accuracy (Low coherence: M = 79.83% correct, SD = 7.86%; High coherence: M = 90.5%, SD = 5.43%; t(22) = 7.3, p<.001) and shorter response times (Low coherence: M = 1176 ms, SD = 149 ms; High coherence: M = 989 ms, SD = 149 ms; t(22)=-11.65, p<.001) (Fig. 1b). Neither accuracy nor RT varied systematically with the direction of motion (upwards vs. downwards; Accuracy: t(22)=1.88; p=0.07; RT: t(22)=0.07, p=0.95). The location of the sensory evidence did influence behavior, such that subjects tended to respond faster and more accurately when the stimulus was presented on the right side of the screen (Accuracy: t(22)=-2.4, p=.02; RT: t(22)=4.23, p<.001).

#### iEEG

We first identified contacts that exhibited task-responsive activity, measured as a significant (p<.01, uncorrected) increase in high-frequency activity (HFA; 70-170 Hz) amplitude during the decision period (see Methods). Of the 3243 contacts included in the analysis (Fig. 1d), we found 963 contacts that showed a significant HFA response following the transition from incoherent to coherent motion (Fig. 2a-c). These task responsive sites were distributed widely across the brain (Fig. 2a, c) and initial inspection of their activity indicated heterogeneity in time courses and alignment to evidence onset vs. response execution (Fig. 2d). All subsequent analyses of the neural data were performed on the subset of contacts identified at this step.

**Figure 2.**
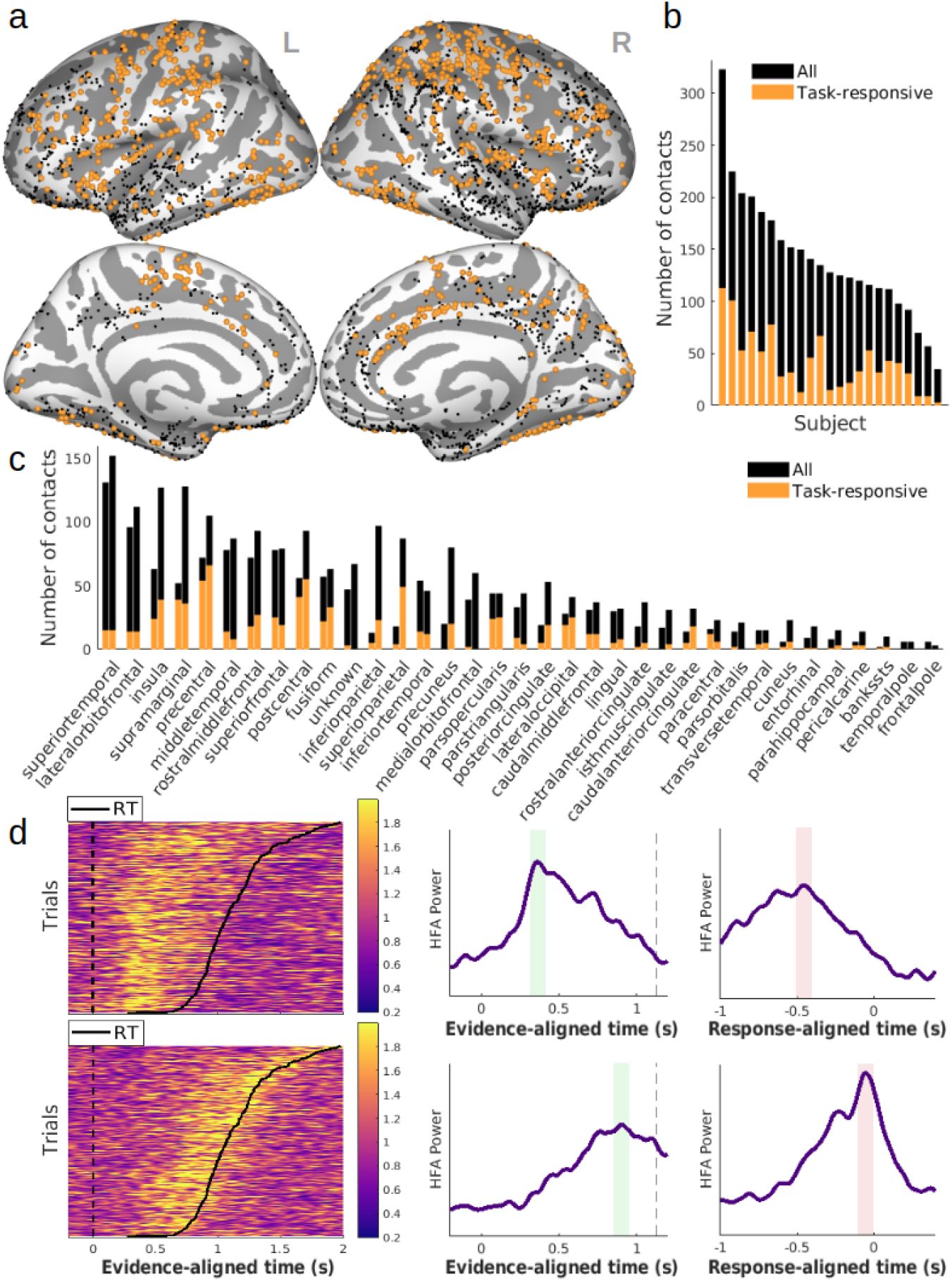
**a.** Spatial distribution of task-responsive contacts in the manual response task (i.e., exhibiting a significant increase in HFA following the onset of sensory evidence). **b**. Number of contacts per subject (black: all contacts included in the analysis, orange: task-responsive sites). **c.** Distribution of contacts across cortical regions. Region labels reflect gyral-based neuroanatomical parcellation^33^. **d.** Activity patterns from two example contacts with task-responsive HFA, illustrating distinct alignment to onset of sensory evidence vs. motor response (top: superior parietal; bottom: precentral gyrus). Color scale represents HFA amplitude. Left: single trial HFA as a function of time. Right: trial-averaged HFA as a function of time. Dashed line marks the median response time. Green and red shaded areas show average peak HFA amplitude in evidence-aligned and response-aligned data, respectively.

We proceeded to categorize task-responsive contacts based on their activity profile, focusing on how HFA at these sites varied with stimulus properties and motor effector. Specifically, for each contact, we performed a 2×2×2×2 ANOVA to assess whether and how the HFA activity depended on sensory evidence strength, spatial location of the evidence, the side of the motor response, and the alignment to evidence onset vs. the time of response (see Methods). Only trials associated with correct responses were included in this analysis.

Our main goal was to localize neural activity consistent with effector-independent evidence accumulation, as distinct from low-level sensory encoding and sensorimotor signals. To this end, we first looked for activity consistent with the latter two categories.

We reasoned that activity relating to expressing choice (i.e., involving motor planning, selection, and/or execution) would exhibit effector-selectivity at, or near, the time of the motor response. Therefore, we searched for contacts that showed a significant effect of Motor Effector (left vs. right hand) on HFA amplitude, with stronger activity for responses made with the contralateral effector relative to the recording site (thresholded at p<.05, uncorrected). Additionally, we expected that effector-selective activity would be temporally aligned to, and reach peak magnitude at or near the time of motor response, in line with previous reports from monkey electrophysiology and human scalp EEG^5,24^ - importantly, this would lead to larger average peak amplitudes in the response-aligned vs. the evidence-aligned trial-averaged data, due to inherent trial-to-trial variability in response times. Thus, we set as an additional criterion a significant effect of Temporal Alignment (i.e., stronger HFA at the time of response-aligned vs. evidence-aligned peak; p<.05, uncorrected). We identified a total of 218 contacts meeting these criteria, which, as expected, were primarily located in sensorimotor regions of the cortex (Fig. 3a, top panel).

**Figure 3.**
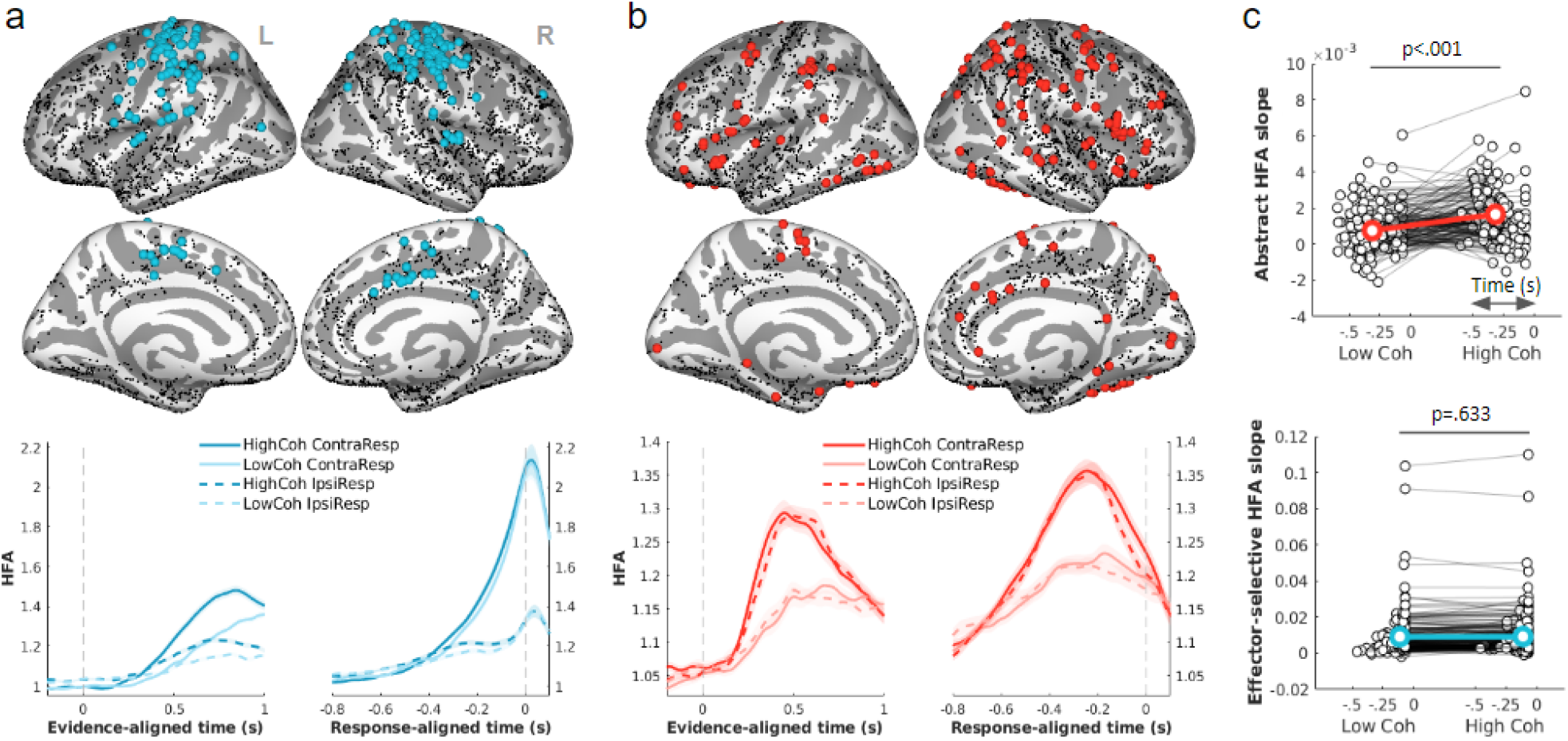
Spatiotemporal profile of HFA consistent with: **a.** effector-selective, mainly response-aligned activity and **b.** abstract (effector-independent) activity. **Top:** Spatial distribution (colored dots: sites meeting criteria for activity profile of interest; black dots: all sites included in the analysis). Electrodes have been projected onto the Freesurfer^32^ average inflated brain (light gray: gyri; dark gray: sulci). **Bottom:** Temporal profile of activity, separated by strength of sensory evidence (High vs. Low motion coherence) and effector laterality (contralateral vs. ipsilateral hand response), averaged across contacts. Shaded areas represent standard error of the mean across contacts. **c.** Modulation of response-aligned buildup rate by motion coherence for abstract accumulator candidate sites (top) and effector-selective sites (bottom). Each pair of data points represents an individual contact. The slopes of the HFA buildup were computed by fitting a straight line to the trial-averaged trace, separately for each motion coherence level, in a 200 ms time window ending at the response-aligned peak HFA amplitude (with peak amplitude identified within the −600 to 30 ms time interval relative to motor response); x axes show the central points of the time windows used to estimate the HFA buildup rate at each contact. The spatial distribution of response-aligned HFA peak times can be seen in Supplementary Fig. 2. For electrodes categorized as effector-selective, only trials associated with the contralateral motor response were included in this analysis.

We next searched for activity consistent with transient, low-level sensory encoding. Non-invasive EEG has characterized transient signals appearing relatively early which, similarly to evidence accumulation signals, scale with the strength of sensory evidence but are distinct in being closely aligned to evidence onset and lateralized according to the hemifield in which evidence appears^28,34^. Our behavioral paradigm involving two simultaneous stimuli (one in each hemifield) allowed us to selectively search for activity meeting these characteristics. Thus, we looked for contacts that simultaneously exhibited 1) significant modulation by Evidence Location (with the expectation of stronger activation when the coherent motion was presented on the side contralateral to the contact; p<.05, uncorrected); 2) significant positive modulation by Evidence Strength in the evidence-aligned HFA (p<.05, uncorrected); 3) a significant effect of Temporal Alignment (p<.05, uncorrected), with greater mean amplitude when aligned to the onset of the evidence than to the motor response; and 4) no significant modulation by left/right Motor Effector (p>.05, uncorrected). We found only 4 contacts that met these criteria, located in the intraparietal sulcus, superior frontal sulcus, and posterior portions of the inferior and middle temporal gyri (Supplementary Fig. 1). As these contacts were so few and mostly situated in sparsely sampled brain areas, we did not further characterize the activity in this category.

Lastly, we looked for sites that exhibited properties similar to the abstract (motor-independent) accumulation activity observed in scalp EEG. To achieve this, we isolated contacts that simultaneously showed 1) significant modulation by the Evidence Strength (p<.05, uncorrected), whereby stronger motion coherence at the evidence onset aligned peak was associated with higher HFA amplitude (HFA _EV-ALIGN, HIGH-COH_ > HFA _EV-ALIGN, LOW-COH_), 2) independence from the limb used to express choice, defined as the absence of a significant main effect of Motor Effector (p>.05, uncorrected), and 3) independence from the spatial location of the sensory evidence, defined as the absence of a significant main effect of Evidence Location (p>.05, uncorrected). As we do not yet know whether and how the neural representations of the choice alternatives (upward and downward motion directions) are intermingled vs. spatially separated in the cortex, much less whether iEEG could distinguish any such spatial separation, our approach did not include direction selectivity as a criterion. Additionally, while we used temporal alignment to evidence onset and motor response as selection criteria when isolating effector- and target-selective contacts, respectively, we did not do so when localizing candidates for abstract evidence accumulation. Existing literature would suggest that the culmination of the evidence accumulation process tends to be more tightly aligned to the time of motor execution (i.e., with choice commitment being shortly followed by a response), which would result in the average signal amplitude showing a higher peak in the response-aligned compared to the evidence-aligned data. However, this is not always observed to be the case (e.g., see ^35^) and is likely to vary across tasks depending on the proportion of RT variance that is driven by evidence accumulation vs. post-commitment motor execution processes. We therefore did not impose any assumptions regarding the relative degree of alignment to evidence onset vs. response execution when isolating these abstract signals. Under these criteria, we identified 161 contacts distributed broadly across prefrontal, parietal, ventral temporal, and insular regions (Fig. 3b, top panel). Activity in these sensors grew gradually over time and peaked several hundred milliseconds before response execution (Fig. 3b, bottom panel). As a further validation step we tested for an effect of evidence strength on the build-up rate of response-aligned activity across the 161 contacts (Fig. 3c). We compared build-up rates associated with the two motion coherence conditions by performing one-tailed paired t-tests across contacts, under the prediction of steeper build-up rates for the High coherence condition. Indeed, we found that across contacts in this class, activity buildup was reliably modulated by evidence strength (p<.001; Fig. 3c, top panel). This was in contrast to activity in the effector-selective category, where this effect was not significant (p=.633) across contacts (see discussion).

#### Relationship between decision-related HFA and behavior

Having located candidate abstract evidence accumulation signals on the basis of sensitivity to key task components, we sought to further investigate their relationship with choice performance, employing criteria that did not inform their initial identification. Non-invasive EEG recording studies have highlighted a dissociation in the way that effector-selective and abstract evidence accumulation signals relate to choice behavior. Whereas effector-selective signals reach a fixed amplitude at response irrespective of accuracy or RT, the amplitude of abstract cumulative evidence signals prior to choice reduces alongside accuracy with increasing RT when strict response deadlines are imposed^27,30^. This feature has been shown to accord with the operation of evidence-independent ‘urgency’ signals at the motor level that progressively reduce the cumulative evidence required to trigger action^27,30,36,37^. To establish whether a similar dissociation was evident in the present data, we first calculated conditional accuracy functions (CAF) by dividing trials into six RT bins for each subject and computing the proportion of correct responses in each bin. With the exception of the fastest RT bin, accuracy tended to decrease monotonically with increasing RT (5×2 repeated-measures ANOVA on the proportion of correct responses across the five fastest RT bins within each motion coherence condition, F_RT_(4,76)=13.2, p<.001; Fig. 4a, top panel).

**Figure 4.**
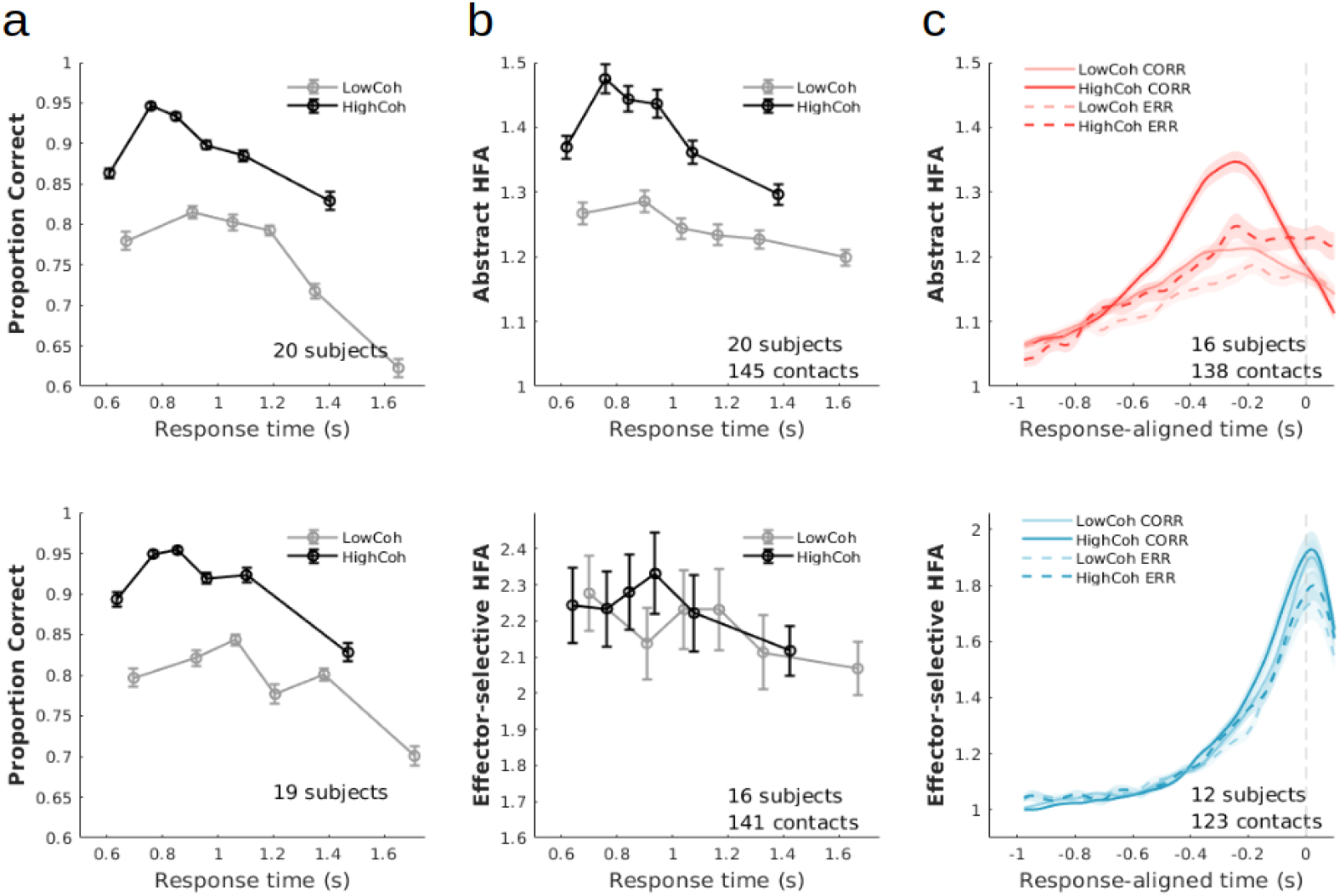
Relationship of HFA with decision-related behavior. Top and bottom panels represent data corresponding to the effector-independent (abstract) and effector-selective electrodes, respectively. **a.** Conditional accuracy function: choice accuracy as a function of response time bins, separated by the strength of sensory evidence (i.e., motion coherence). The y axis shows the average accuracy across subjects (weighted by the number of contacts that contributed to each subject’s data). **b.** HFA at the response-aligned peak, as a function of response time bins. The y axis shows the average HFA across all contacts in a given electrode category. Only trials associated with correct choices were used for this analysis. **c.** Response-aligned average HFA traces, separated by choice accuracy and evidence strength. All analyses in **a-c** reflect data from subjects with a minimum of 5 trials available per bin. For electrodes categorized as effector-selective, only trials associated with the contralateral motor response were included in the analysis.

As predicted, for abstract accumulator candidates, the relationship between RT and peak HFA amplitude in the response-aligned signal (Fig. 4b, top panel) mimicked the pattern observed in the conditional accuracy function, with activity decreasing as a function of RT, except for the fastest RT bin (5×2 repeated measures ANOVA on HFA amplitude: F_RT_(4,576)=36.18, p<.001). We verified this similarity by computing Pearson’s correlations between the RT-binned pre-response HFA and corresponding choice accuracy, separately for each channel. To assess whether the average correlation across channels in this category was significantly higher than chance, we generated a null distribution by computing the average correlation coefficient after shuffling the order of HFA bins, and repeating this procedure 5000 times. We found that the two patterns of modulation by RT were significantly correlated (mean R=.376, p=.01). This pattern is in line with an abstract decision-related evidence accumulation signal where slower, less accurate decisions are made based on smaller amounts of cumulative evidence. In contrast, in sites characterized by effector-selective activity (Fig. 4, bottom panels), correlation with the CAF was not significant (mean R=.05, p=.54; Fig. 4b, bottom panel) and was lower than that observed for HFA from abstract accumulator candidates (t(284)=9.88, p<.001). Similarly, the decrease in HFA amplitude with slower RTs was significantly weaker at these sites (2×2 mixed ANOVA with factors Contact Category and Motion Coherence on the rate of HFA decrease with RT, i.e., slopes obtained from fitting a straight line to the average HFA associated with the five slowest RT bins; F_CATEGORY_(1, 284)=21.46, p<.001).

We also observed higher amplitudes for correct responses than for errors in the HFA from abstract accumulator candidates, in line with errors being based on less cumulative evidence (2×2 repeated-measures ANOVA across contacts, with factors Motion Coherence and Accuracy; F_ACC_(1, 137)=56.46, p<.001; Fig. 4c, top panel). Interestingly, this effect was also present in the effector-selective HFA (F_ACC_(1,122)=83.06, p<.001), although given that activity at these locations tended to peak at or even after motor response, it is possible that the higher amplitudes observed here for correct choices might be related to fluctuations in motor execution (e.g., vigor of response).

### Vocal response task

Although we made efforts to isolate potential generators of abstract evidence accumulation from effector-selective activity by excluding sites that exhibited limb selectivity, it is nevertheless conceivable that activity at some of these locations could be movement-dependent in a way that is not selective to the limb used to respond. Thus, to provide additional support for motor independence in our candidate sites, we examined recordings from a subset of participants who performed an alternative version of the task (Fig. 1a), where requirements for modality and timing of response were modified (Methods). Specifically, subjects were asked to report their choice vocally, and to do so only after a forced random delay (1000-2000 ms) marked by a visual cue.

#### Behavior

Subjects (n=11) responded correctly on 88.28% of the trials (SD = 8.05%). Choice accuracy was not significantly different from the manual response task (t(10)=.89, p=.39). Similarly to the manual response task, behavioral performance was modulated by the strength of sensory evidence, with subjects responding correctly more frequently on trials with strong sensory evidence (Low coherence: M = 81.22% correct, SD = 12.36%; High coherence: M = 95.35%, SD = 4.83%; t(10)=4.86, p<.001) (Fig. 1c).

#### iEEG

We inspected data from the 11 subjects who took part in both the motor and vocal response tasks. Of the 161 sites identified as candidates for effector-independent evidence accumulation in the manual response task, 82 were also recorded from during the vocal response task (Fig. 5a, Supplementary Fig. 3). We sought to investigate whether activity at these locations continued to display signature characteristics of evidence accumulation despite the change in motor requirements. Indeed, we continued to see an increase in HFA after the onset of sensory evidence, as well as modulation of amplitude by the strength of sensory evidence in 42 out of 82 contacts (evaluated within a 100ms time window centered on the evidence-aligned peak; thresholded at p<.05, uncorrected; see Methods). For completion, the modulation of HFA by evidence strength across all task-responsive electrodes common between the two tasks can be visualized in Supplementary Fig. 4.

**Figure 5.**
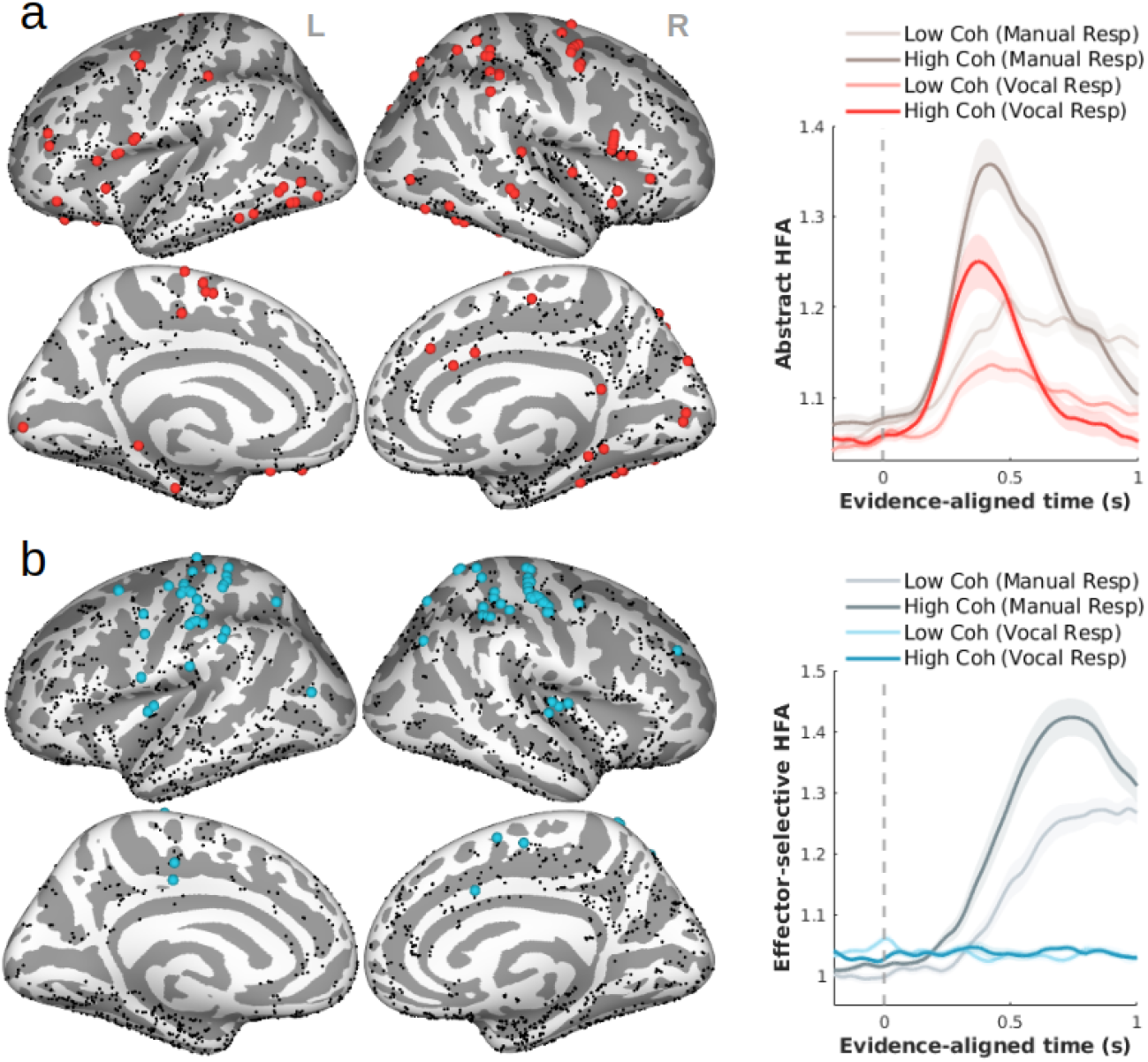
HFA during the delayed vocal response task in sites previously categorized as **a.** Abstract (i.e., independent of hand response laterality) and **b.** Selective for effector laterality. *Left panels:* Black dots show contact coverage during the vocal response task. Colored dots mark contacts where neural recordings were made during both the manual and vocal response tasks, and which met categorization criteria in the analysis of the manual response task data. *Right panels:* Temporal profile of average HFA, shown separately for data recorded during the speeded manual response (gray) vs. vocal response task (color), and separated by strength of sensory evidence (High vs. Low motion coherence). Shaded areas represent standard error of the mean across contacts.

We performed the same comparison for sites we previously identified as showing effector-selective activity (Fig. 5b). Neural recordings during the vocal task were available for 79 of the 218 of these contacts. As expected of effector-selective signals, these regions no longer exhibited a neural response to the onset of the sensory evidence or amplitude modulated by the strength of the evidence. Activity at only 2 out of the 79 sites continued to show modulation by evidence strength in the vocal task (thresholded at p<.05, uncorrected). Overall the predominant absence of task-related responses in this subset of contacts suggests that if activity at these sites supports the decision process, it does so in a manner that is closely tied to the motor action associated with the choice.

## Discussion

Localizing the brain regions involved in the formation of movement-independent perceptual decisions is a long-standing challenge in decision research. By exploiting the spatiotemporal precision of intracranial EEG, our study sought to provide a more extensive spatial mapping of abstract decision-related signals in the human brain than was previously available. We identified a broadly distributed network where activity exhibited characteristics consistent with abstract representations of evolving perceptual decisions. Specifically, we observed a gradual build-up following the onset of sensory evidence, at a rate that scaled with the quality of the evidence and reflected both subjects’ choice accuracy and response times. Crucially, these signals were insensitive to the evidence location and laterality of the motor effector. Key to our approach was the ability to further validate the independence of these decision-related signals from movement requirements using an independent data set, showing that activity continued to trace cumulative evidence irrespective of whether participants responded via a speeded manual button push or delayed vocalization.

Our results are unique in that they identify abstract, evidence-dependent decision dynamics in the human brain which have previously only been observed using non-invasive methods characterized by high temporal but relatively poor spatial precision^17,23,24^. Our approach allowed us to not only capture the focal activity that might have been missed with scalp EEG, but to sample from multiple regions across the brain from our large sample of participants. The spatial distribution of the abstract accumulation dynamics observed in our data included prefrontal, parietal, inferior temporal, and anterior insular regions. Not surprisingly, several of these regions have been previously implicated in effector-independent decision processes, by work involving human neuroimaging (prefrontal cortex, PFC^16^, anterior insula^13,14^, inferior temporal^38^), and single unit recordings in humans (parietal^39^) and non-human primates (PFC^10^, parietal^9^), though none of these studies identified all of these activations at once.

The profile of activity at the abstract accumulator candidate sites identified here was consistent with human scalp EEG signals previously shown to exhibit evidence accumulation dynamics in an effector-independent manner^25,29,30,35,40^. A question arising from this observation is whether our data highlight potential neural generators of such CPP-like signals. It should first be noted that the local dynamics responsible for voltage fluctuations measured with scalp EEG are likely distinct from those underlying the HFA measurements obtained from intracranial recordings. Specifically, while event-related potentials reflect relatively slow fluctuations in the broadband signal arising primarily from dendritic postsynaptic potentials^41^, intracranially recorded HFA has been shown to correlate, at least in part, with neuronal spiking activity^42,43^. Thus, to the extent that activation in any given area would tend to give rise to both spiking and postsynaptic activity, it is plausible that one or more of the areas identified here contribute to the scalp-measured CPP, though the degree of the overlap remains speculative for now. Future work employing simultaneous recordings of scalp and intracranial EEG and/or source modeling may provide further insight into this question.

The relationship of the neural signal with stimulus properties and behavior in the period just before overt choice commitment has important implications for interpreting the quantity represented at these locations. Specifically, we showed that, in keeping with observations of the scalp-recorded CPP, the peak amplitude of the neural response scaled with the strength of the sensory evidence, as well as with response speed and decision accuracy. This pattern differs from what is typically observed in movement-selective regions, where activity reaches a fixed threshold prior to choice irrespective of stimulus properties or behavior^3,44^. Insight into the origin of this difference comes from work investigating decision dynamics under varying speed pressure conditions. Specifically, it has been proposed that in the context of speeded perceptual decisions (i.e., where a choice deadline is imposed), an additional time-dependent “urgency” signal may contribute to elevating the decision variable to an action-triggering boundary in areas of the brain responsible for planning and preparing the appropriate movement^36,37,45,27^, such that smaller amounts of cumulative evidence are necessary for committing to a choice as time elapses. According to this view then, abstract evidence accumulation signals like the CPP and the iEEG signals isolated in the present study reflect the cumulative evidence informing the choice at the time of commitment which varies as a function of urgency. In contrast to the abstract accumulator candidates, activity at the effector-selective sites was more in line with a boundary-crossing process, with activity reaching similar amplitudes regardless of the amount of sensory evidence. However, unlike movement-selective evidence accumulators in the non-human primate brain, where response-aligned activity also reflects the steeper buildup rate for easier choices, we did not, on average, see such an effect in this group of contacts. This could be because our effector-selective criteria would identify not only signals related to motor preparation, to which there is a continuous flow from evidence accumulation, but also signals related to motor execution and/or somatosensory feedback, which would be more tightly response-aligned, stereotyped, and possibly dominant over motor preparation signals. Indeed, non-invasive neurophysiology work has shown that motor preparation signals tend to be reflected more reliably in lower frequency bands whereas higher frequency responses tend to be more response-aligned and stereotyped, similar to what we found using intracranially recorded HFA^46^.

Interestingly, the activity at our abstract accumulator candidates differed from that of CPP-like EEG signals in terms of its timing relative to choice commitment. Average HFA in our study peaked considerably earlier (i.e., 200-300 ms prior to motor response) than the CPP which is typically observed to peak at, or just prior to the response. One possible explanation is that the scalp-recorded CPP may contain a mixture of both abstract evidence accumulation and motor-related signals (planning/execution). As we excluded sources of effector-selective activity in our analysis, our data may reflect a more precise temporal profile of abstract decision formation, whereby the culmination of abstract evidence accumulation precedes that of motor-related processes. Alternatively, the time between choice commitment and motor execution might have been longer for our patient sample due to limited training and time on the task compared to the typical experiments performed in healthy volunteers.

It remains to be determined whether the sites identified here form a homogenous network actively involved in evidence accumulation or whether they have unique contributions, and which, if any, play a causal role in the decision formation process, as opposed to merely tracking the cumulative evidence. Future investigations able to perturb neural activity during evidence accumulation (e.g., intracranial electrical stimulation^47^) may help answer these questions. Also worth considering is that, although we recorded from a large participant sample with substantial bilateral coverage of all cortical lobes, our study is subject to a limitation inherent to all intracranial studies, namely heterogeneous electrode coverage across patients due to electrode locations being determined solely based on clinical purposes. It is therefore possible that our study could have missed decision-related signals originating in regions not sampled from, for instance subcortical structures such as the basal ganglia^48,49^, which have previously been implicated in evidence accumulation.

## Methods

### Participants

We recorded data from neurosurgical patients suffering from drug-resistant epilepsy and undergoing intracranial EEG (iEEG) monitoring at the North Shore University Hospital (Manhasset, NY 11030, USA), aimed at locating seizure foci and informing treatment. The study was approved by the institutional review board of the Feinstein Institutes for Medical Research. All participants provided written informed consent in accordance with the Declaration of Helsinki.

23 participants (11 female, age range 18-57, M = 33.9 years) took part in the speeded manual response version of the random dot task. Of these, 11 (4 female, age range 18-57, M = 36.7 years) additionally performed a delayed vocal response version of the task. To control for learning effects across the two tasks, we alternated the order of the two tasks, such that 5 out of 11 subjects performed the vocal version first, while the remaining 6 subjects performed the manual version first.

### Stimuli and tasks

Stimuli were presented at bedside using the Psychophysics Toolbox version 3 for MATLAB (Mathworks, Natick, MA). Stimuli consisted of two simultaneous random dot kinematograms (white dots on black background) presented within circular apertures that were placed symmetrically on the left and right of a central fixation dot (Fig. 1a). At the onset of the dot stimuli, motion on both sides was incoherent. After a random delay (1.4-1.8 s, uniformly distributed), motion on one unpredictable side of the fixation became coherent towards one of two possible directions (up or down). Coherent motion lasted for 2 s. The transition from incoherent to coherent motion served to minimize the extent to which sensory evoked potentials to visual changes might obscure decision-related activity. Subjects were asked to determine the direction of the coherent motion, irrespective of the side it was presented on. Offset of the motion stimulus was followed by performance feedback for that trial: “Correct”, “Wrong” or “Too early”/”No response?” on trials where a response was made before the onset of coherent motion, or missed the response deadline, respectively. Feedback was displayed for .4 s and was followed by an inter-trial interval of 1 s (blank screen).

Difficulty of the random dot stimuli was determined by the proportion of dots moving coherently in one direction (i.e., motion coherence) and was calibrated separately for each subject during a practice session prior to the main experiment. We used two motion coherence levels, adjusted to span perceptual peri-threshold (~75-80% correct). Motion coherence for the difficult condition was adjusted to half of the value of the easy condition. The average proportions of coherently moving dots for the Low and High coherence conditions across the 23 subjects were .28 and .56, respectively. All stimulus conditions (evidence presentation side, motion coherence, and motion direction) were equally and randomly distributed across trials.

#### Speeded manual response task

On each trial, participants reported their choice by making a button press with their left or right hand using a mouse. The two possible motion direction choices were each assigned to one motor effector (left or right limb), and this mapping was maintained throughout the entire experiment. Choice-effector mapping was counterbalanced across subjects (10/23 subjects responded with right hand to report upward motion). Subjects performed between 3-6 blocks of 64 trials each (average number of trials per subject: 256). Motor responses were allowed only during the time window where coherent motion was present. Trials where responses were given outside of this time range were excluded from further analysis (mean proportion of missed trials across subjects = 8.87%; SD = 7.92%).

#### Delayed vocal response task

The vocal version of the task was identical to the motor version, with two exceptions: 1) subjects were instructed to wait through a random delay (1-2 s, uniformly distributed) for a visual “go” cue before reporting their choice, which was marked by a change in the fixation color from white to green; 2) choices were reported vocally by naming the direction of the dots out loud (e.g., “Up” or “Down”).

### iEEG data acquisition

Patients were implanted with either depth (2- or 1.3-mm platinum cylinders with 4.4- or 2.2-mm center to center distance and a diameter of 0.8 mm; PMT corporation, Chanassen, MN) and/or subdural strip/grid electrodes (2- or 3-mm platinum disks with 4- or 10-mm center to center distance; PMT corporation, Chanassen, MN). The placement of the electrodes was determined exclusively based on clinical considerations. Intracranial data were recorded using either an XLTEK Quantum Amplifier (Natus Medical Inc., San Carlos, CA) at 1 kHz, or a Tucker-Davis Technologies (Alachua, FL) acquisition module, at 3 kHz. The iEEG signal was referenced online to a reference electrode located either subdurally, subdermally, or in the white matter, which was determined upon visual inspection of the signal quality, at the bedside. Transistor-transistor logic (TTL) pulses triggered by the stimulus presentation software at events of interest were recorded simultaneously with the iEEG data, which subsequently served to align intracranial recordings to task-related events of interest.

### Electrode localization

Electrode localization and visualization was performed using the iELVis toolbox^50^ for MATLAB. Prior to electrode implantation, a T1-weighted 1-mm isometric structural magnetic resonance imaging (MRI) scan was acquired for each subject, on a Siemens 3T MRI scanner. After electrode implantation, a computerized tomography (CT) scan and a T1-weighted MRI scan were acquired. The post-implantation CT and MRI scans were co-registered to the preoperative MRI scan using FSL’s BET^51^ and FLIRT^52,53^ algorithms. Co-registration of these two scans allowed us to minimize localization error caused by potential brain shift and to visualize the CT scan on top of the preoperative MRI. Semi-manual identification of contact artifacts on the co-registered CT was done in BioImageSuite^54^. Volumetric information from the preoperative T1 scan was obtained using the “recon-all” command in Freesurfer^32^ v6.0.0.

### Data preprocessing

Electrodes located in seizure onset zones and those that displayed abnormal epileptiform discharges, as assessed by the clinical epileptology team, were excluded from further analysis. iEEG time series were then visually inspected for signal quality, and any electrodes containing systematic artifacts or interictal spikes were also discarded. We also excluded contacts located fully in white matter tissue, using the proximal tissue density (PTD) measurement^55^ as a criterion. The PTD represents the normalized difference between gray and white matter voxels surrounding the contact centroid (where min/max values of −1/1 reflect tissue composition of only white and only gray matter, respectively). To maintain a conservative approach to electrode exclusion, we removed only those for which PTD<.-9. Overall, a total of 3243 contacts across all subjects were considered for further analysis (Fig. 1a). Data were resampled to 512 Hz. Additionally we applied notch-filters at 60 Hz, 120 Hz, and 180 Hz. Lastly, data were re-referenced to the average of all electrodes that passed the quality check. We epoched the data from −3 to 3s relative to the onset of the sensory evidence (i.e., coherent motion). Trials containing strong transient artifacts were manually removed based on visual inspection.

### HFA analysis

To extract high frequency activity (HFA), we bandpass filtered the preprocessed signal in non-overlapping 10Hz bands from 70 to 170 Hz (using Matlab “filtfilt”; 4th order butterworth). We then applied the Hilbert transform to the filtered signal and extracted the instantaneous amplitude from the resulting analytic signal. The instantaneous amplitude was normalized by division to the average of the time series, and squared to extract power.

### Channel/trial selection

All analyses were performed on trials associated with correct responses, unless otherwise specified. Task-responsive sites were defined as those that exhibited an overall increase in HFA during the “decision period” (i.e., the time interval starting 200 ms after evidence onset and ending at the subject-specific median response time), relative to a pre-evidence baseline period (−450 ms to −50 ms prior to onset of the coherent motion), as assessed with a one-tailed t-test on time-averaged HFA (α=.01, one-tailed t-test).

### ANOVA

For data acquired during performance of the manual response task, we examined the relationship between HFA and three task-related variables of interest: 1) the strength of sensory evidence of the random dot stimulus (i.e., High vs. Low motion coherence), 2) the motor effector used to report choice (i.e., Ipsilateral vs. Contralateral limb, relative to the recording site), and 3) the spatial location of the sensory evidence (Ipsilateral vs. Contralateral side of the visual fixation). We simultaneously assessed these effects in the evidence-aligned and response-aligned data. Specifically, for each contact, we extracted the mean HFA within a 100ms window centered on the peak of trial-averaged activity, separately for evidence- and response-aligned data. For evidence-aligned data, selection of the average peak was made from the interval starting 200ms after evidence onset and up to 100 ms after the subject’s median RT. For response-aligned data, the peak search interval started 600ms before, and ended 30 ms after motor response. We conducted a 2×2×2×2 mixed model ANOVA on these measurements, with between-unit factors: Evidence Strength (High/Low), Motor Effector (Ipsilateral/Contralateral Limb), Evidence Location (Ipsilateral/Contralateral Side), and within-unit factor Time of Measurement (Evidence-/Response-aligned).

For data acquired during the vocal response task, we assessed modulation by evidence strength with a 2×2 ANOVA on evidence-aligned HFA, with factors Evidence Strength and Evidence Location. The 100ms time window of analysis for each electrode was selected under the same parameters as above, using subject-specific median RTs from the manual response task as estimates for the end of the decision period.

## Supplementary material

**Supplementary Fig. 1.**
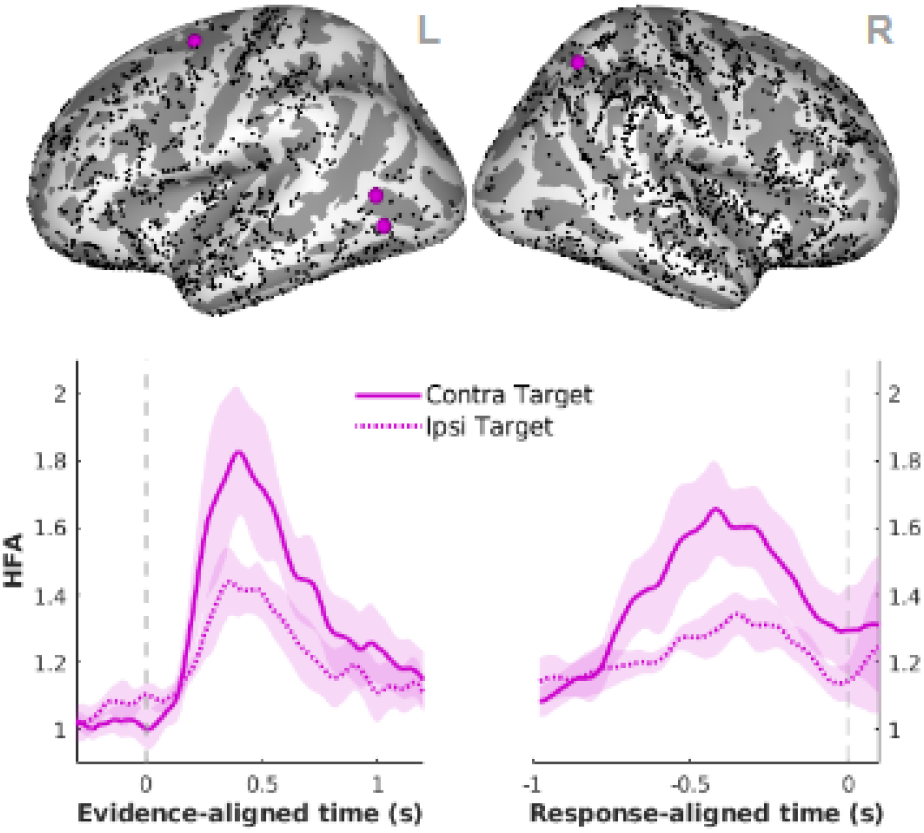
Spatiotemporal profile of HFA consistent with transient low-level sensory encoding. *Top:* Spatial distribution (black dots: all sites included in the analysis). *Bottom:* Temporal profile of activity, separated by strength of sensory evidence (High vs. Low motion coherence) and spatial location of the evidence (Contralateral vs. Ipsilateral side), averaged across contacts. Shaded areas represent standard error of the mean across contacts.

**Supplementary Fig. 2.**
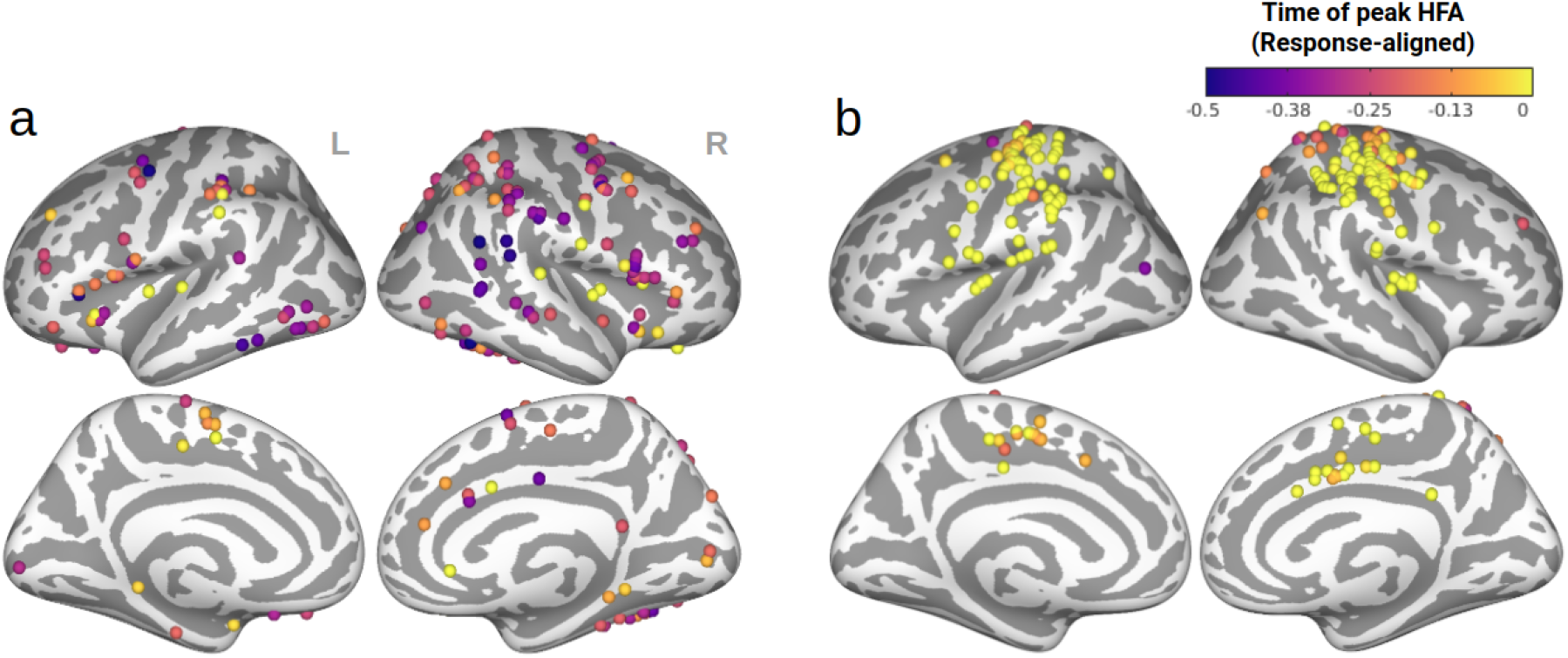
Time of peak amplitude in response-aligned, trial-averaged HFA across electrodes categorized as abstract (i.e., effector-independent) accumulator candidates (**a**) and effector-selective (**b**).

**Supplementary Fig. 3.**
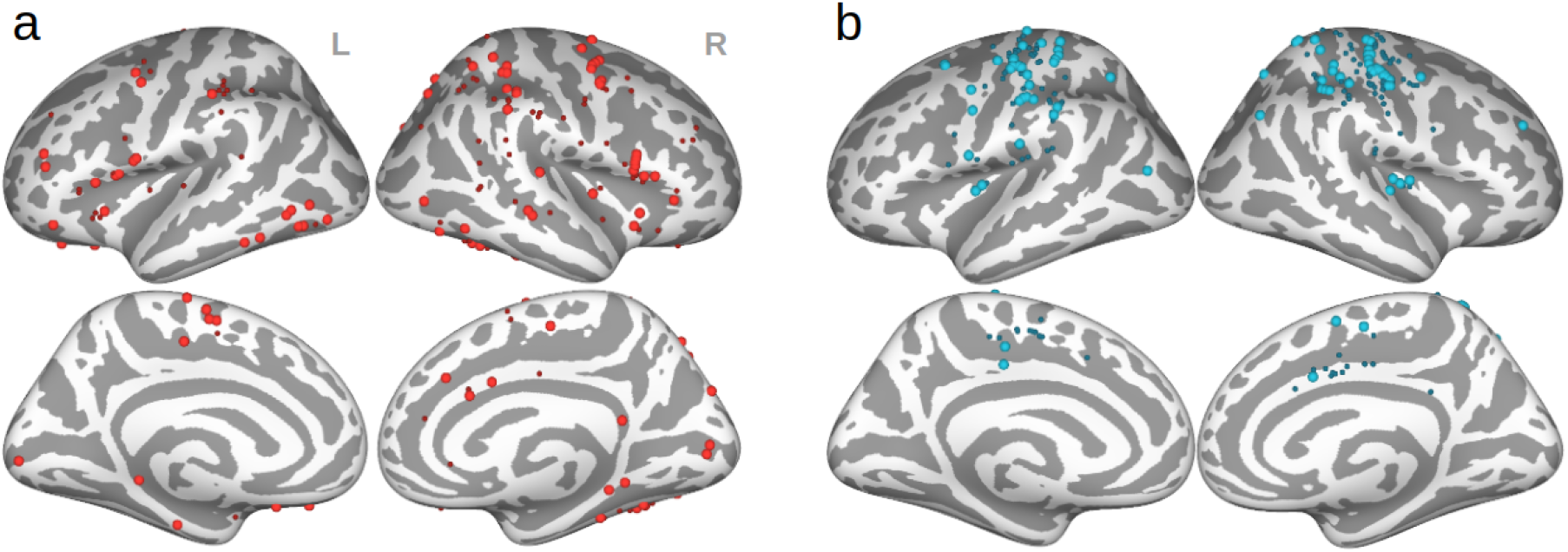
Electrodes categorized as abstract (i.e., effector-independent) accumulator candidates (**a**) and effector-selective (**b**) based on analysis of the manual response task data. Large dots mark the subset of these electrodes which were also recorded from during the vocal response task.

**Supplementary Fig. 4.**
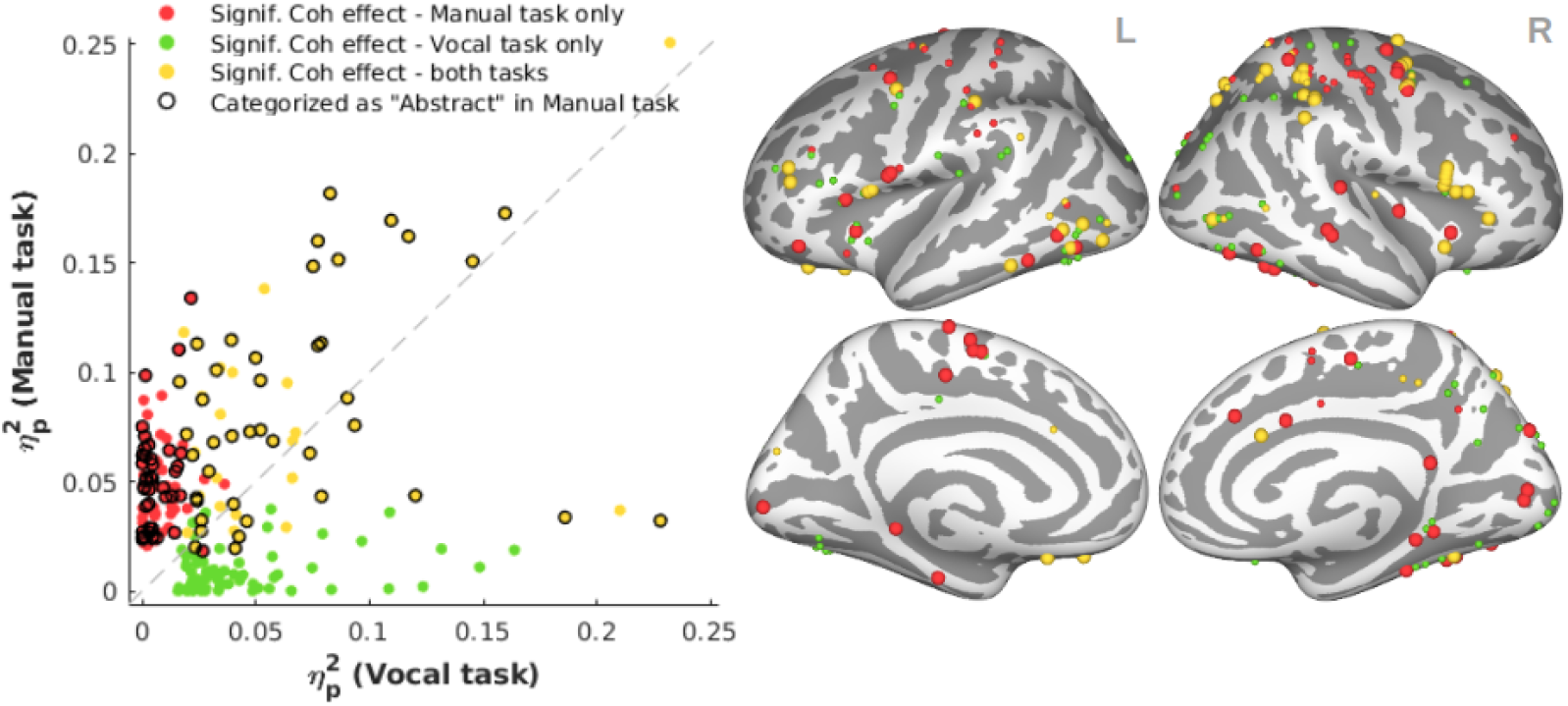
Modulation of HFA by evidence strength in contacts common between the two behavioral tasks. Displayed here are only the task-responsive contacts showing a significant effect of evidence strength (High>Low) in the evidence-aligned signal in at least one of the tasks (thresholded at p<.05, uncorrected; see ANOVA in Methods). **a.** Effect sizes (partial eta squared, η_p_^2^) reflecting the magnitude of the modulation by evidence strength during the vocal (*x* axis) and manual (*y* axis) response tasks. Data points represent individual contacts. **b.** Spatial distribution of evidence strength modulation (large dots: contacts categorized as abstract accumulator candidates during the manual-response task).

